# A general strategy to red-shift green fluorescent protein based biosensors

**DOI:** 10.1101/2020.04.09.034561

**Authors:** Shen Zhang, Hui-wang Ai

## Abstract

Compared to green fluorescent protein (GFP) based biosensors, red fluorescent protein (RFP) based biosensors are inherently advantageous because of reduced phototoxicity, decreased autofluorescence, and enhanced tissue penetration. However, there is a limited choice of RFP-based biosensors and development of each biosensor requires significant effort. Herein, we describe a general and convenient method which uses the genetically encoded amino acid, 3-aminotyrosine (aY), to convert GFPs and GFP-based biosensors into red.

---

Fluorescent protein (FP) based biosensors are indispensable research tools.^1,2^ Despite that RFP-based biosensors are increasingly reported, the emission of most existing FP-based biosensors falls in the green or yellow spectral region.^2^ Their spectral overlap hinders and sometimes precludes multiplexing. Moreover, RFP-based biosensors are expected to reduce phototoxicity and autofluorescence and increase photon penetration and imaging depth. However, existing RFP-based biosensors often suffer from small dynamic range, mislocalization, and undesired photoconversion.^2^ Thus, in reality RFP-based biosensors are repeatedly outperformed by their green fluorescent counterparts. In this context, there is a pressing need to expand RFP-based biosensors for not only a broader range of analytical targets but also enhanced properties.

Although several green-to-red, photoconvertible FPs have been reported and widely used,^1^ there is limited success to directly convert a typical GFP to an RFP. In one study, mutagenesis of GFP resulted in an “R10-3” mutant containing a mixture of green and red fluorescent chromophores.^3^ A few other studies reported green-to-red photoconversion under anaerobic conditions,^4,5^ under extensive ultraviolet or blue irradiation with high GFP expression levels,^6,7^ or in the presence of electron receptors.^8,9^ Nevertheless, these approaches usually lead to incomplete photoconversion; there is no reliable evidence that they can be generalized for green-to-red conversion of diverse GFP-like proteins and their derived biosensors.

We here present a general and convenient method, which uses genetic code expansion^10^ to introduce a noncanonical amino acid (ncAA), aY (**Fig. 1a**), to the chromophores of GFP-like proteins and biosensors for spontaneous and efficient green-to-red conversion.

**Figure 1.**
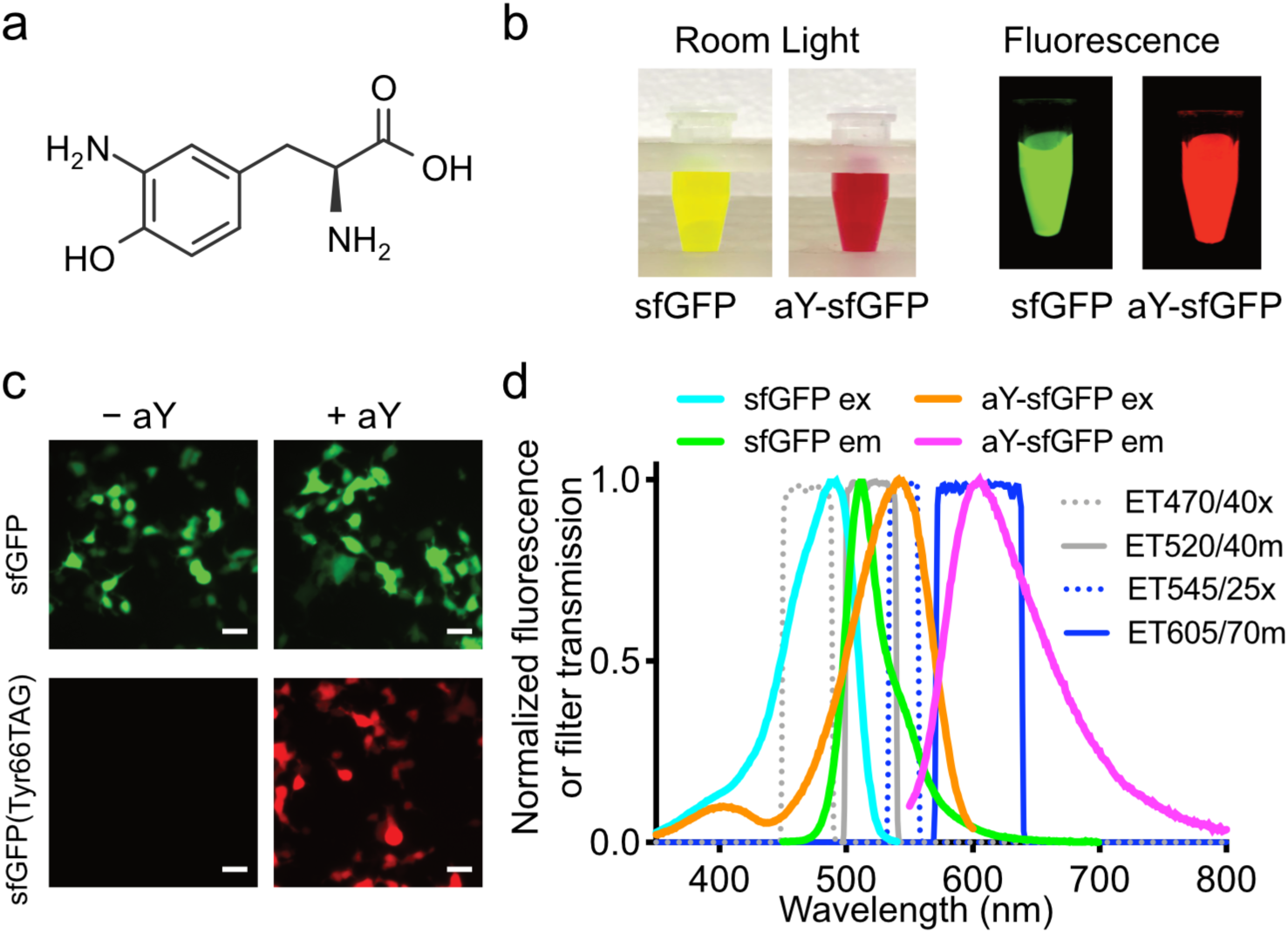
Green-to-red conversion of sfGFP by 3-aminotyrosine (aY). (**a**) Chemical structure of aY. (**b**) Imaging of sfGFP and aY-sfGFP proteins prepared from *E. coli*. (**c**) Microscopic imaging of HEK 293T cells co-transfected with pMAH-*Ec*aYRS and pcDNA3-sfGFP or pcDNA3-sfGFP(Tyr66TAG), showing red fluorescence of aY-sfGFP in mammalian cells. Scale bar: 20 µm. (**d**) Fluorescence excitation and emission spectra of sfGFP (cyan and green) and aY-sfGFP (orange and magenta), suggesting that sfGFP and aY-sfGFP can be sequentially imaged with little crosstalk using appropriate filters. Transmission spectra for excitation and emission filters designated with Chroma Technology part numbers are also plotted (gray and blue). The experiments in panel **b-d** were repeated three times with similar results.

By expressing orthogonal tRNA and aminoacyl-tRNA synthetase (aaRS) in live cells, diverse ncAAs can be site-specifically introduced into proteins in response to nonsense or four-base codons.^10^ To genetically introduce aY into proteins in *E. coli*, we first examined a reported *Methanococcus jannaschii* tyrosyl-tRNA synthetase (*Mj*TyrRS) mutant but observed high cross-reactivity with Tyr.^11^ Next, we examined another *Mj*TyrRS mutant, which was initially engineered for the genetic encoding of 3-iodotyrosine.^12^ We computationally docked aY to the active site of this enzyme and introduced randomization.^13^ We identified a promising *Mj*TyrRS mutant (designated *Mj*aYRS. Next, we expressed a superfolder GFP (sfGFP)^16^ mutant with an amber codon introduced to its residue 66 in the presence of *Mj*aYRS and the corresponding amber suppression tRNA. We observed strong fluorescence and obtained the full-length protein in good yield (∼ 6.7 mg per liter of culture) when aY was supplemented. We further characterized the resultant protein with electrospray ionization mass spectrometry (ESI-MS) and confirmed the incorporation of aY to sfGFP.

To genetically encode aY in mammalian cells, we took *E. coli* tyrosyl-tRNA synthetase (*Ec*TyrRS) as our starting point. We first introduced a D265R mutation to increase anticodon recognition by the aaRS,^14^ and further inserted a tyrosine editing domain into *Ec*TyrRS to reduce its reactivity with tyrosine.^15^ We next created 30 *Ec*TyrRS variants with mutations at residues 37 and 195 within the amino acid-binding pocket. Screening of these mutants for amber suppression of pcDNA3-EGFP(Tyr39TAG) identified an aaRS variant (*a*.*k*.*a. Ec*aYRS). In the presence of aY, *Ec*aYRS, and the corresponding tRNA, full-length EGFP was generated, as confirmed by both fluorescence microscopy and ESI-MS.

The color of aY-sfGFP (sfGFP^16^ with aY introduced to its chromophore-forming residue 66) from *E. coli* was red under both room light and green excitation, and it showed nearly no residual green fluorescence (**Fig. 1b**). To test whether this phenomenon is species-specific, we expressed aY-sfGFP in human embryonic kidneys (HEK) 293T cells and then observed spontaneous, red fluorescence (**Fig. 1c**). The excitation and emission maxima of aY-sfGFP were red-shifted from those of sfGFP by 56 and 95 nm, respectively, suggesting that it may be possible to pair aY-sfGFP with GFPs or GFP-based biosensors for sequential, dual-color imaging using common fluorescence microscope setups (**Fig. 1d**).

The chromophore of GFP is spontaneously formed through cyclization, dehydration, and oxidation of an internal tripeptide motif, while the chromophores of common RFPs differ from GFP in that additional, self-catalyzed oxidation occurs to expand chromophore conjugation via a hydrolyzable *N*-acylimine substitution.^17^ Because aY is sensitive to oxidation, we hypothesize that aY-sfGFP spontaneously forms an RFP-like chromophore through additional oxidation. Although the detailed mechanism remains to be explored, the formation of a hydrolyzable chromophore is supported by backbone cleavage of aY-sfGFP under denaturing conditions. Furthermore, the absorbance of aY-sfGFP decreases at ∼ 540 nm with concurrent increase at ∼ 390 nm as pH drops from neutral to acidic, suggesting the existence of an anionic chromophore in aY-sfGFP at neutral conditions.

Next, we introduced aY to the chromophores of several FPs containing Tyr-derived chromophores, including teal fluorescent mTFP1,^18^ yellow fluorescent Citrine,^19^ a circularly permuted sfGFP (cpsGFP),^20^ and a circularly permuted yellow FP (cpYFP).^21^ Red-shifted excitation and emission were observed for all of them, indicating that the aY-induced green-to-red conversion is neither protein-specific nor topology-specific.

After confirming that aY can red-shift various Tyr-derived chromophores, we used aY to modify a number of biosensors based on circularly permuted green, teal, or yellow FPs, including G-GECO1 (a Ca^2+^ sensor),^22^ ZnGreen1 (a Zn^2+^ sensor),^23^ iGluSnFR (a glutamate sensor),^24^ iGABASnFR (a GABA sensor),^25^ dLight1.2 (a dopamine sensor),^26^ SoNar (a NAD^+^/NADH sensor),^27^ iNAP1 (a NADPH sensor),^28^ PercevalHR (an ATP sensor),^29^ and iATPSnFR1.1 (another ATP sensor).^30^ We observed red-shifted fluorescence for all test biosensors (**Table 1**). Moreover, the converted biosensors largely retained the dynamic range and responsiveness of their green fluorescent predecessors, as shown in our validation experiments with purified proteins, cultured mammalian cell lines, and/or primary neurons.

**Table 1.** Properties of genetically encoded biosensors and their aY-modified variants.

| Biosensor | Analytical target | Excitation (nm) | Emission (nm) | Dynamic Range ($\Delta F/F_{\min}$)<sup>a</sup> | References |
| --- | --- | --- | --- | --- | --- |
| G-GECO1 | Ca<sup>2+</sup> | 497 | 512 | 1,983% | Zhao et al.<sup>22</sup> |
| aY-G-GECO1 | Ca<sup>2+</sup> | 528 | 594 | 1,566% | *This work* |
| ZnGreen1 | Zn<sup>2+</sup> | 474 | 500 | 2,531% | Chen et al.<sup>23</sup> |
| aY-ZnGreen1 | Zn<sup>2+</sup> | 520 | 585 | 915% | *This work* |
| iGluSnFR | glutamate | 488 | 510 | 257% | Marvin et al.<sup>24</sup> |
| aY-iGluSnFR | glutamate | 528 | 600 | 270% | *This work* |
| iGABASnFR | GABA | 502 | 513 | 143% | Marvin et al.<sup>25</sup> |
| aY-GABASnFR | GABA | 541 | 602 | 92% | *This work* |
| dLight1.2 | dopamine | 496 | 513 | 194% | Patriarchi et al.<sup>26</sup> |
| aY-dLight1.2 | dopamine | 545 | 603 | 117% | *This work* |
| SoNar | NAD<sup>+</sup>/NADH | 497 | 512 | 257% | Zhao et al.<sup>27</sup> |
| aY-SoNar | NAD<sup>+</sup>/NADH | 544 | 604 | 426% | *This work* |
| iNap1 | NADPH | 497 | 513 | 82% | Tao et al.<sup>28</sup> |
| aY-iNap1 | NADPH | 545 | 604 | 614% | *This work* |
| PercevalHR | ATP | 498 | 513 | 185% | Tantama et al.<sup>29</sup> |
| aY-PercevalHR | ATP | 545 | 604 | 120% | *This work* |
| iATPSnFR1.1 | ATP | 493 | 513 | 106% | Lobas et al.<sup>30</sup> |
| aY-iATPSnFR1.1 | ATP | 541 | 608 | 75% | *This work* |

<sup>a</sup> Defined as intensimetric changes ( $\Delta F/F_{\min}$ ) around emission peaks in the presence of 480 and 540 nm excitation for green and aY-modified, red fluorescent biosensors, respectively.

In summary, we genetically encoded aY in both *E. coli* and mammalian cells, and developed a general method based on the introduction of aY to the chromophores of GFP-like proteins and biosensors for spontaneous and efficient green-to-red conversion. This approach does not require photoconversion or other special experimental conditions and is compatible with a variety of FPs and FP-based biosensors in *E. coli*, mammalian cell lines, and primary cells. Little optimization is needed as dynamic range and responsiveness are largely retained in the converted biosensors. Therefore, this method can be used to quickly expand the toolkit of RFP-based biosensors.

## ACKNOWLEDGEMENTS

We thank R. Campbell, L. Looger, B. Khakh, G. Yellen, and P. Schultz for plasmids, S. Chen for MIN6 cells, and other members of the Ai laboratory for discussion and assistance with experiments. We thank Drs. Wei Ren and Ao Ji for early exploration of this project. Research reported in this publication was supported in part by the University of Virginia and the National Institutes of Health under awards R01GM118675, R01GM129291, and U01CA230817.

## AUTHOR CONTRIBUTIONS

H.A. conceived and supervised the project. S.Z. performed all experiments. H.A. and S.Z. analyzed the data and prepared the manuscript.

